# Adding noise to Markov cohort state-transition models

**DOI:** 10.1101/635177

**Authors:** Rowan Iskandar

## Abstract

Following its introduction over thirty years ago, the Markov cohort model state-transition has been used extensively to model population trajectories over time (Markov trace) in decision modeling and cost-effectiveness studies. We recently showed that a cohort model represents the average of a continuous-time stochastic process on a multidimensional integer lattice governed by a master equation (ME), which represents the time-evolution of the probability function of an integer-valued random vector. Leveraging this theoretical connection, this study introduces an alternative modeling method using a stochastic differential equation (SDE) approach which captures not only the mean behavior but also the variance of the process. The SDE method constitutes the time-evolution of a random vector of population counts. We show the derivation of an SDE model from first principles by postulating the expected change in the distribution of population across states in a small time step. We then describe an algorithm to construct an SDE and solve the SDE via simulation for use in practice. We show the applications of SDE in two case studies. The first example demonstrates that the population trajectories, and their mean and variance, from the SDE and other commonly-used methods match. The second example shows that users can readily apply the SDE method in their existing works without the need for additional inputs beyond those required for constructing a conventional cohort model. In addition, the second example demonstrates that the SDE model is superior to a microsimulation model in terms of computational speed. In summary, an SDE model provides an alternative modeling framework which includes information on variance, can accommodate for time-varying parameters, and is computationally less expensive than microsimulation for a typical modeling problem in cost-effectiveness and decision analyses.

## 1 Introduction

Decision models have been used in various applications from health technology assessment to clinical guideline development. In decision modeling, a Markov cohort state-transition model is often used to simulate the prognosis of a patient or a group of patients following an intervention. Beck and Pauker [1] introduced the modeling method over 30 years ago with the aim to provide a simple tool for prognostic modeling and for practical use in medical decision making. In principle, a Markov cohort state-transition model [1–3] is a recursive algebraic formula that calculates the *average* number of individuals in each state (*state-configuration*) in the next time step as the sum of current state-configuration and the change in the state-configuration during the time step. This calculation produces a deterministic quantity, i.e. the average of the process, and thus does not completely specify a stochastic representation of the modeled system.

We recently explicated the stochastic process underlying a cohort model, defined as a continuous-time integer-valued stochastic process, in the context where population counts are of interest.[3] The stochastic process is governed by a well-known time-evolution equation of a probability function, known as a master equation. Given this knowledge of the evolution equation associated with a cohort model, we can identify the relationship between a cohort model and other known methodologies in stochastic processes literature.[4] In particular, if we relaxed the whole-number (integer) constraint on the state space to allow for fractional (non-integer) numbers, a cohort model can be immediately extended to represent a change in the *random* state-configuration by adding the variance of the process to the average of the process. This representation of a change in a random vector as the sum of its average and noise is known as the stochastic differential equation (SDE) model.[5] Information on the variance of a process is useful for quantifying the inherent variability of the population trajectories[6] and the extent of accuracy in the mean prediction as well as for deciding whether it is worth collecting more information about the decision problem.[7]

This paper aims to introduce an SDE model as an alternative modeling framework by capitulating the relationship between the stochastic process underlying a cohort model and the SDE. For this paper, we adopt a continuous-time perspective, i.e. for a discrete-time process, a corresponding continuous-time process exists (*embeddability-assumption*). In addition, we assume that the process of interest is Markovian, i.e. knowledge of the current state conveys all information that is relevant for forecasting future states. This paper is organized into three main parts. First, we formulate a stochastic cohort model under two representations. We first reintroduce the stochastic process underlying a Markov cohort state-transition model in the form of a master equation. Then, we postulate an SDE representation of the stochastic cohort process which captures the average and the variance of the process. Secondly, we provide a step-by-step guidance for formulating an SDE model given a state-space in the modeling problem, a set of transition rules, and the intensity of each transition. Lastly, we demonstrate two applications of an SDE approach in decision modeling. The first application shows the equivalence between existing methods[3] and an SDE. In the second application, we develop an SDE model for a recently published decision model evaluating the effect of chemotherapy-induced toxicity among breast cancer patients.[8] In this example, we demonstrate the computational efficiency of an SDE model vs. a microsimulation approach in a setting where the transition rates are time-variant. Throughout this exposition, we try to strike a balance between mathematical rigor and accessibility to practitioners.

## 2 Stochastic cohort model

A stochastic cohort model where each individual follows a Markov chain on a finite set of mutually exclusive and completely exhaustive health states: 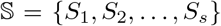, is defined as a stochastic process, {**N**(*t*)}_*t*≥*t*_0__ where *s* is the number of states and *t* is time (*t* ≥ *t*_0_ for an initial time *t*_0_). **N**(*t*) represents a random vector of population counts (*state-configuration*):

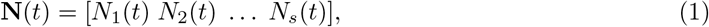

in which 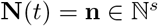 (where 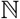 is the set of non-negative integers or counting numbers) with the probability of observing a particular state-configuration, 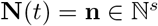 at time *t* or *P* (**N**(*t*) = **n**) = *P*(**n**, *t*). In this section, we first present the master equation which governs the stochastic process underlying a cohort model: a stochastic process on 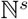. [3] Then, we introduce an SDE formulation of the stochastic cohort model as a stochastic process on ℝ^*s*^ (where ℝ is the set of real numbers).

### 2.1 Master equation

An individual transitioning from *S_k_* to *S_l_* is denoted by the transition *R_i_* where *k*, *l* ∈ {1 …, *s*} and *i* ∈ {1 …, *r*} (*r* is the total number of transitions and *s*^2^ is the maximum number of possible transitions or *r* ≤ *s*^2^). Since the occurrence of each transition alters the state-configuration, each *R_i_* is associated with a vector representing the change in the current state-configuration, *δ_i_*, and is defined as *δ_i_* = *e_l_* − *e_k_* where *e_j_* is the *j*-th column of an identity matrix of size *s*. We associate each transition with a *propensity function*: *ν_i_*(**n**, *t*) = *c_i_*(*t*)*h_i_*(**n**), where *c_i_*(*t*) is the transition rate for transition *R_i_*, and **N**(*t*) = **n**. The function *h_i_* (**n**) determines the number of ways *R_i_* can occur. In the case where an interaction between individuals in two or more different states has no effect, *h_i_* (**n**) is equal to the number of at-risk individuals, e.g. for *S_k_* to *S_l_* transition, *h_i_*(**n**) = *n_k_*. A model for an infectious disease will have more terms in *h_i_* for capturing the rate of contacts between individuals in different disease states. The propensity function denotes the rate of a (any) transition occurring in continuous time as a function of the state-configuration and rate constant. The probability of observing *one particular* transition *R_i_* in a small time interval *τ* is assumed to be linear in time: *c_i_*(*t*)*τ* + *o*(*τ*), where *o*(*τ*) represents other terms that decay to zero faster than *τ*. Therefore, the probability of a (any) transition from *S_k_* to *S_l_* in time step *τ* is given by *n_k_c_i_*(*t*)*τ* + *o*(*τ*).[3] A transition *R_i_* occurring with a probability *ν_i_* (**n**, *t*)*τ* in a time step *τ* alters the current state **N**(*t*) = **n** to:

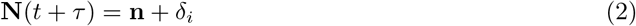

The master equation is obtained by enumerating all possible transitions contributing to a change in the probability of observing a particular state-configuration in an infinitesimal time step [3] and is given by:

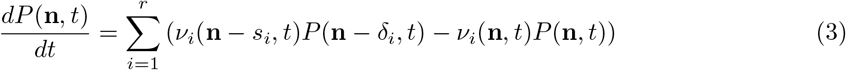

The terms, *ν_i_* (**n**, *t*) *P*(**n**, *t*), represent the probability which is transferred from one to another state-configuration as a result of a transition *R_i_*. In principle, a master equation expresses the change per unit of time of the probability of observing a state-configuration **n** as the sum of two opposite effects: the probability flow into and out of **n**, and exactly represents the dynamics of the stochastic process underlying a cohort model.

### 2.2 Stochastic differential equation

In this section, we seek an alternative representation of a stochastic cohort model. Our aim is to approximate a Master equation by assuming that the change in the state-configuration is allowed to be fractions or non-integer, i.e. **N**(*t*) has a continuous sample path or *N_i_*(*t* + *τ*) − *N_i_*(*t*) → 0 as *τ* → 0 for *i* ∈ {1,2,…, *s*}. In addition, instead working with the evolution equation for the *P* (**n**, *t*), we consider the evolution of a random vector, **N**(*t*). Our approach of deriving an SDE model is a heuristics one.

Given a state space 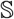 and a set of transitions *R_i_* where *i* ∈ {1 …, *r* } (as in Section 2.1), the time evolution of a random vector, **N**(*t*), can be determined by enumerating all possible changes to **N**(*t*) in a small time step, *τ*, which is governed by the transition type (*R_i_*), its associated change vector (*δ_i_*), and its associated probabilities (*ν_i_*(**n**, *t*)*τ*). From Equation 2, we write the change in the state-configuration in *τ* time step as Δ**N**(*t*) = **N**(*t* + *τ*) − **N**(*t*). The 1 × *s* vector of the expected change in the state-configurations at time *t* (*drift* vector) is given by:

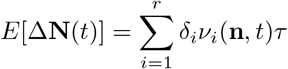

or in matrix form:

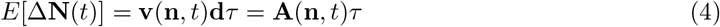

where **v**(**n**, *t*) is a 1 × *r* vector of propensities where the *i*-th column corresponds to the propensity function for the transition *i* (*ν_i_* (**n**, *t*)), **d** is an *r* × s matrix of *δ_s_* where the *i*-th row corresponds to the vector of changes associated with the transition *i*, and **A**(**n**, *t*) is a 1 × *s* vector in which the *k*-th element corresponds to the rate of change in the number of individuals in state *S_k_*. The **A**(**n**, *t*) is essentially the right-hand side of the ordinary differential equation-equivalence to a cohort model.[3, 9] Equation 4 is the average of the change in current state-configuration across possible transitions weighted by their probabilities of occurring. Similarly the covariance matrix of the change in the state-configurations in *τ* step (*diffusion* matrix) can be calculated via the definition of the covariance matrix for a random vector:

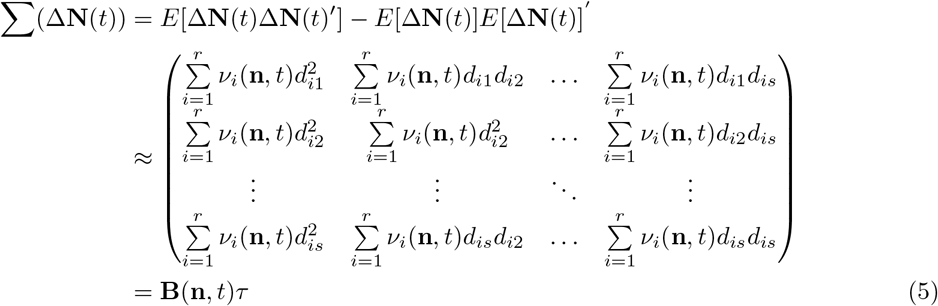

if the terms of order higher than *τ* vanish. Let *ξ*(*t*) be a *s*-dimensional standard Wiener process, i.e. *ξ*(*t*) follows a standard normal distribution or 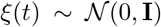, where **I** is an *s* × *s* identity matrix. Using a Normal approximation: 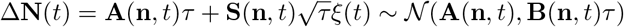, the evolution of a stochastic cohort model can be represented by the following updating formula:

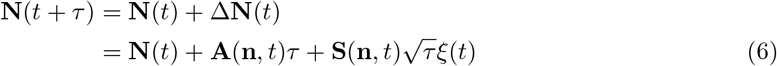

where 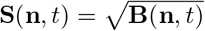 (see Appendix A for a more detailed explanation). If *τ* approaches 0, then we have 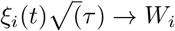 and arrive at the *Itô* SDE:

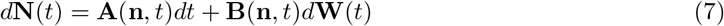

where **W**(*t*) be a *s*-dimensional Wiener process. A more theoretical discussion on stochastic differential equations can be found in [5] and [10].

## 3 Developing an SDE

In this section, we describe an algorithm for developing an SDE model for applications in decision modeling. The algorithm involves the following steps:

**S1** determine the state space 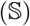, all possible transitions (*R_i_*) among the states in 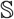, and their corresponding propensities, *ν_i_*, for *i* ∈ {1, …, *r*};
**S2** construct the 1 × *r* vector of propensities, **v**(**n**, *t*), and the *r* × *s* matrix of changes, **d**;
**S3** calculate the drift vector (Equation 4) and the diffusion matrix (Equation 5);
**S4** choose a numerical method for approximately solving the SDE by generating a sample path (also known as *Markov trace*): 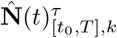 where *τ*, [*t*_0_, *T*], and *k*, refer to the numerical method time step, the time interval of analysis, and the index of the samples, respectively;
**S5** and estimate a quantity of interest as a function of the sample path(s), i.e., 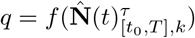, where: *q* is the quantity of interest and *f* is a mapping from the sample paths to the quantity of interest.

The first two steps are identical to specifying a deterministic cohort model. To calculate the drift vector and the diffusion matrix, we use Equations 4 and 5, respectively. Standard methods to diagonalize a positive semi-definite matrix [11] can be utilized to calculate the square root of the diffusion matrix.

For solving an SDE model, a numerical method, such as the Euler-Maruyama method [12], is often used since in most applications, particularly for models with time-varying parameters (e.g. age-specific probability of dying to other causes), an analytical solution to an SDE would not be feasible. A typical numerical method is based on the idea of dividing the time interval of interest, e.g. [*t*_0_, *T*], to *n* discrete time points, which may not necessarily be uniformly-spaced: *t*_0_ = *τ*_0_ < *τ*_1_ < *τ*_2_ < … < *τ*_*n*−1_ < *τ_n_* = *T*, and estimating the values of **N**(*t*) at these time points sequentially. For example, the Euler-Maruyama approximation, 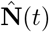, of an SDE model, *d***N**(*t*), is given by the following recursive equation:

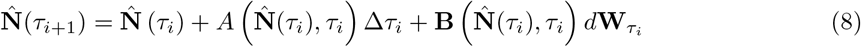

where Δ*τ_i_* = *τ*_*i*+1_ − *τ_i_* and *d***W**_*τi*_ = **W**(*τ*_*i*+1_) − **W**(*τ_i_*) denotes the increment of the Wiener process in the time interval [*τ_i_*, *τ*_*i*+1_] and is represented by independent Gaussian random variables with zero mean and variance of Δ*τ_i_* = *τ*_*i*+1_ − *τ_i_*. In practice, the *d***W**_τ*i*_ is approximated by 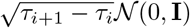. An application of the numerical method will yield a sample path: 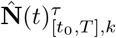.

Quantities of interest, such as survival, prevalence or lifetime costs, and their statistics (mean and variance) can be estimated empirically by generating a *sufficiently* large number of sample paths. For example, if *q* represents the life expectancy of a cohort, then its estimator, 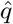, can be estimated by simulating *L* number of sample paths:

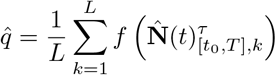

where the mapping *f* is defined as the ratio of the total sum of individuals across non-death states in [*t*_0_, *T*] to the initial population size. In general, the accuracy of the estimator depends on the choice of the Euler-Maruyama’s size, *τ_s_*, and the number of sample paths, *L*, which are problem-dependent.

## 4 Applications of SDE

We conduct two case studies to demonstrate: (1) the equivalence of an SDE model to other commonly-used methods in decision modeling, and (2) how to develop a stochastic cohort model using an SDE approach. In the second case study, we also compare the computational time between the SDE and microsimulation approach since both approaches produce estimate of the variance and easily accommodate for time-varying rates. In both case studies, the models are coded in R [13] and Julia [14] as open-source programs have emerged as the lingua franca for practitioners in decision modeling. [15, 16] The comparison of both methods’ code execution time is conducted in Julia. The codes for the second example are available under a GNU GPL license and can be found at https://github.com/rowaniskandar/SDE.

### 4.1 Case study: 4-state Markov model

For the first example, we emulate the example in [3] to show the connection between SDE and other modeling methods: (1) a cohort model with *τ* = 1-year, (2) a microsimulation model, and (3) an SDE, which is solved with the Euler-Maruyama method and *τ* = 1-year. We consider a generic four-states stochastic process (*s* = 4), i.e. 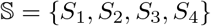 (Figure 1), with rate constants of *c*_1_ = 0.05, *c*_2_ = 0.01, *c*_3_ = 0.001, *c*_4_ = 0.1, *c*_5_ = 0.05, and *c*_6_ = 2. We simulate a cohort of 1000 individuals, who start in *S*_1_, i.e. *N*_1_(0) = 1000, for a 50-years duration. To calculate the mean and the variances of **N**(*t*) based on the SDE model, we generate 10 and 10000 sample paths. The benchmark for comparing variances is based on 10 and 10000 Monte Carlo samples of a *microsimulation population* with *τ* = 1-year. Each microsimulation population is conceived as a realization of 1000 Markov chains (or individual trajectories). For the cohort model, the calculation of the 1-year transition probability matrix from the transition rates is based on the formulas from Welton and Ades. [17] The results of the population trajectories in state S3 are given in Figure (Figure 2). We observe no difference in the trajectories between the empirical estimates from the SDE, cohort model and microsimulation. Also, the estimates of the standard deviations from both SDE and microsimulation approaches are approximately equal (Figure 2).

**Figure 1.**
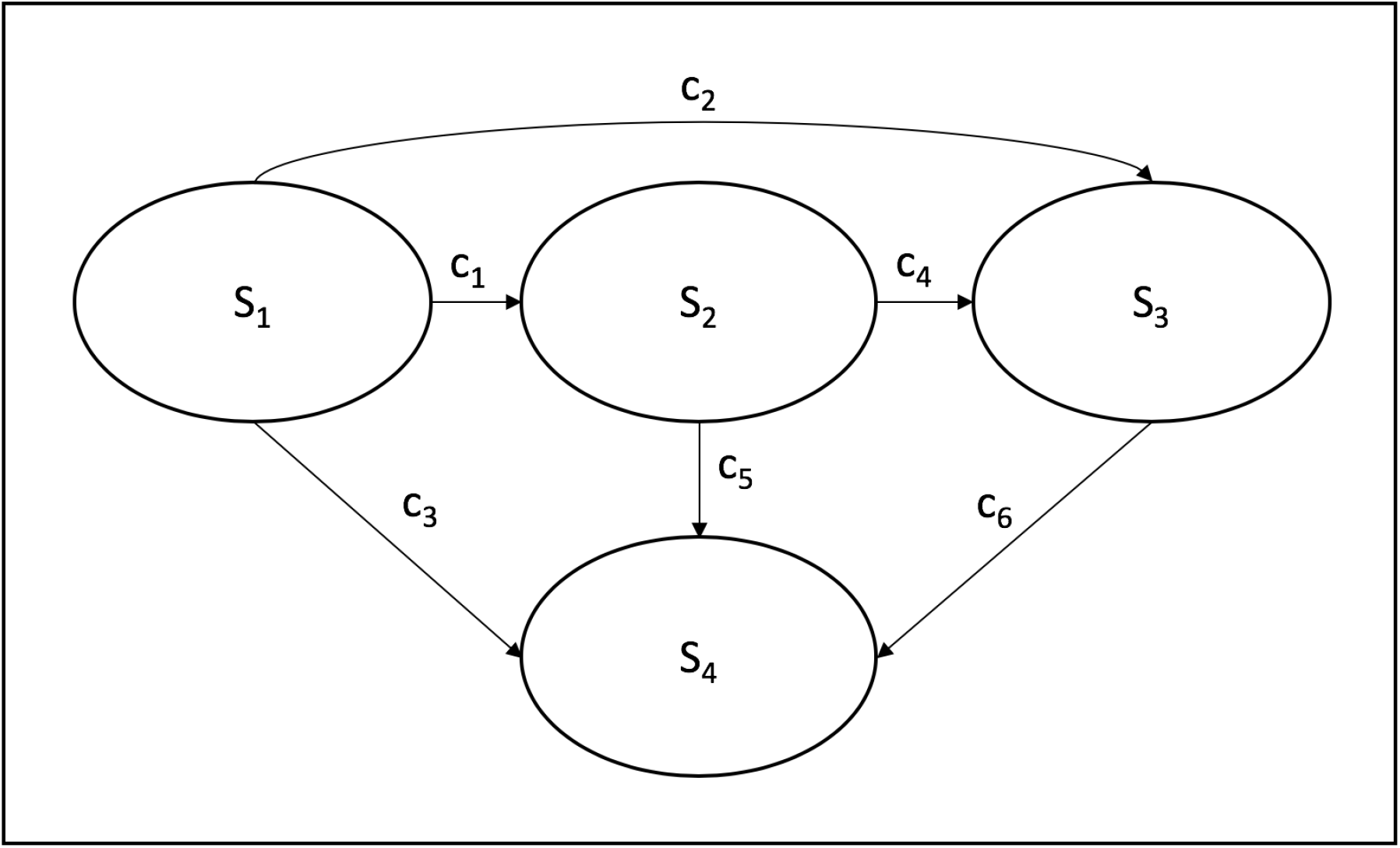
A state-transition model diagram used in case study 1

**Figure 2.**
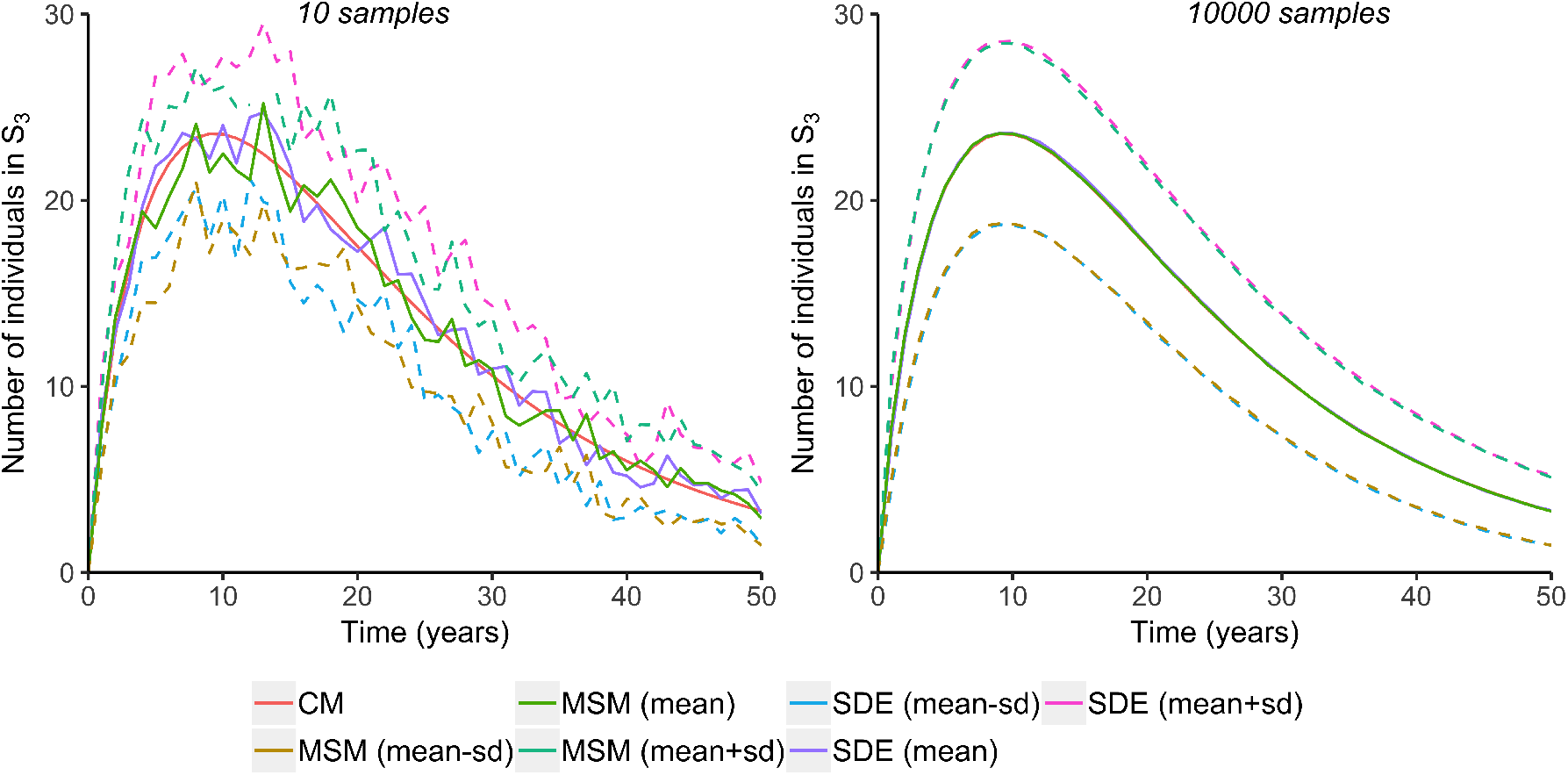
Population trajectories for state S3 in case study 1 by different methods and Monte Carlo sample sizes [CM: cohort model, MSM: microsimulation, SDE: stochastic differential equation, sd: standard deviation. Dashed lines indicate standard deviations and are not applicable for CM. Plots for 10000 samples are on top of each other]

### 4.2 Case study: Breast cancer toxicity model

For the second example, we develop an SDE model for a published Markov decision model which estimates the life expectancy associated with receiving chemotherapy among 55 year-old breast cancer patients.[8] In developing the SDE model, we follow the steps in Section 3.

**S1** The published (deterministic) cohort model has seven states (*s* = 7): healthy (SH), healthy with treatment toxicity (*S_T_*), metastasis (*S_M_*), metastasis with treatment toxicity (*S_TM_*), death due to treatment toxicity, death due to metastasis (*S_DM_*), and death due to other causes (*S_DO_*). There are 12 possible transitions among the states (Table 1).
**S2** We construct the 1 × 12 vector of propensities **v**(**n**, *t*). The elements of the propensity vector **v** is constructed in the following way. For example, an event of developing a metastasis from a healthy state (*S_H_* → *S_M_*) can occur with a propensity equal to the transition rates times the at-risk individuals in *S_H_*: *c_M_N_H_*(*t*). We then specify the 12 × 7 matrix of changes, **d**. For example, the transition *S_H_* → *S_M_* is associated with the vector [−1 0 1 0 0 0 0]. The ordering of transitions (indexing of rows) in **d** should be consistent with the ordering in **v**(**n**, *t*) (indexing of columns).
**S3** Using the formula (Equation 4), we derive the 1 × 7 drift vector:

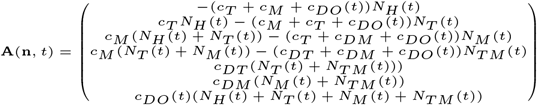 The 7 × 7 diffusion matrix can be similarly constructed by using Equation 5. For example, the first and the fourth columns of **B**(**n**, *t*) are equal to:

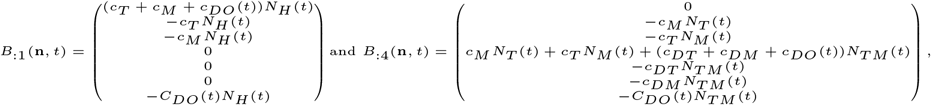

respectively; where each *c*. denotes a transition rate and the subscript corresponds to the (destination) state name. In the model, the rate of death due to other causes varies by age. The description and estimate of each parameter are given in [8].
**S4** Given the drift vector and the diffusion matrix, we apply the Euler-Maruyama method to generate 10000 sample paths of a cohort of 1000 55 year-old women, using a 1-year time step. We use the R package YUIMA [18] and develop a Euler-Maruyama-based solver for Julia.
**S5** From these sample paths, we estimate the life expectancy by summing the number of individuals across non-death health states for all times and divide the sum by the population size.

**Table 1:** Propensities and vectors of changes for breast cancer toxicity model in case study 2 [*S*.: states; *c*.: transition rates; *N*.: number of individuals; ._*H*_: healthy; ._*T*_: treatment toxicity; ._*M*_: metastasis; ._*TM*_: metastasis with treatment toxicity; ._*DT*_: death due to treatment toxicity; ._*DM*_: death due to metastasis; ._*DO*_: death due to other causes]

| Transitions | Propensities (elements of vector $\mathbf{v}$) | Vectors of changes (elements of matrix $\mathbf{d}$) |
| --- | --- | --- |
| $S_H \rightarrow S_T$ | $c_T N_H(t)$ | $[-1 \ 1 \ 0 \ 0 \ 0 \ 0 \ 0]$ |
| $S_H \rightarrow S_M$ | $c_M N_H(t)$ | $[-1 \ 0 \ 1 \ 0 \ 0 \ 0 \ 0]$ |
| $S_H \rightarrow S_{DO}$ | $c_{DO} N_H(t)$ | $[-1 \ 0 \ 0 \ 0 \ 0 \ 0 \ 1]$ |
| $S_T \rightarrow S_M$ | $c_M N_T(t)$ | $[0 \ -1 \ 0 \ 1 \ 0 \ 0 \ 0]$ |
| $S_T \rightarrow S_{DT}$ | $c_{DT} N_T(t)$ | $[0 \ -1 \ 0 \ 0 \ 1 \ 0 \ 0]$ |
| $S_T \rightarrow S_{DO}$ | $c_{DO} N_T(t)$ | $[0 \ -1 \ 0 \ 0 \ 0 \ 0 \ 1]$ |
| $S_M \rightarrow S_{TM}$ | $c_T N_M(t)$ | $[0 \ 0 \ -1 \ 1 \ 0 \ 0 \ 0]$ |
| $S_M \rightarrow S_{DM}$ | $c_{DM} N_M(t)$ | $[0 \ 0 \ -1 \ 0 \ 0 \ 1 \ 0]$ |
| $S_M \rightarrow S_{DO}$ | $c_{DO} N_M(t)$ | $[0 \ 0 \ -1 \ 0 \ 0 \ 0 \ 1]$ |
| $S_{TM} \rightarrow S_{DT}$ | $c_{DT} N_{TM}(t)$ | $[0 \ 0 \ 0 \ -1 \ 1 \ 0 \ 0]$ |
| $S_{TM} \rightarrow S_{DM}$ | $c_{DM} N_{TM}(t)$ | $[0 \ 0 \ 0 \ -1 \ 0 \ 1 \ 0]$ |
| $S_{TM} \rightarrow S_{DO}$ | $c_{DO} N_{TM}(t)$ | $[0 \ 0 \ 0 \ -1 \ 0 \ 0 \ 1]$ |

As a comparison, we also develop a microsimulation model with the following generator matrix:

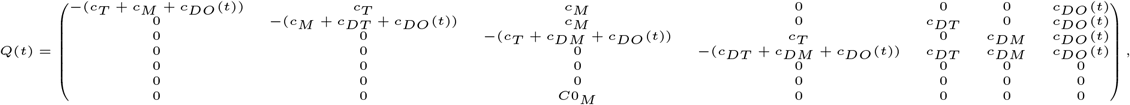

and generate 10000 sample paths with 1-year time step. The life expectancies for the chemotherapy arm are estimated to be 16.9(standard deviation: 0.42) and 17.2(standard deviation: 0.41) for the SDE and microsimulation methods, respectively. Using a benchmarking tool in Julia,[14] the SDE model requires a considerably less computational time than the microsimulation model to generate the 10000 sample paths: 1.5 vs. 51.9 minutes.

## 5 Discussions

In this study, we introduce an SDE method which models not only the average behavior but also the variance of the stochastic process and can readily accommodate for time-varying parameters. In general, one can model the behavior of a stochastic dynamics system by considering either the time-evolution of the random vector (exemplified by SDE) or of the probability function of the random vector (exemplified by master equation). Although the probability function approach potentially provides a way to derive an explicit form of the probability function (i.e. the complete knowledge of the stochastic system) without relying on numerous numerical realizations of the system, the random vector approach is more applicable in the following context. In most applications, an analytical solution to the probability function method is almost inaccessible; thereby rendering numerical methods as the common solution method. In particular, some of the transition rates in most decision models are time-varying, for example: rate of death due to other causes is a time-dependent parameter. In this time-variant setting, practitioners often use Monte Carlo simulation (microsimulation) to estimate decision-relevant quantities from a cohort model. However, this approach, albeit flexible, is computationally expensive. To estimate the variance of the population counts across states, we need to replicate the microsimulation population a number of times *in addition* to the individual Monte Carlo runs within each population, which is a potentially computationally intensive task and is similar to applying uncertainty analysis to a microsimulation model.[19] In particular, if some of the events in the model are rare and/or the expected differences in the effect size among strategies are small, we expect the required number of Monte Carlo runs to be high. Although an SDE model also requires a Monte Carlo solution approach, the sample path of a cohort is generated more efficiently compared to microsimulation: SDE simulates a cohort *in a batch vs. individually*. Indeed, the cohort size would affect the computational speed of a microsimulation but not that of an SDE. The computational efficiency of the SDE vs. microsimulation approach as shown in the breast toxicity model example with time-varying parameter supports the aforementioned assertion.

SDE models have been applied in various fields such as finance[20], infectious disease [21], physics[22], and biology[23]. To date, no study has examined the potential application of an SDE model, as an alternative to a Markov cohort model, in decision modeling. The stochastic process underlying the commonly-used cohort model, defined as a continuous-time discrete-state stochastic process, has only been recently demonstrated.[3] Hence, the theoretical connection between a cohort model and an SDE was unknown prior to [3]. In this study, we provide a guidance to walk users through the process of developing an SDE model. Developers of Markov cohort models should be familiar with the first step which involves: enumerating the possible health states, assigning the allowable transitions to all pairs of states, and specifying the intensity of each transition (which can be time-invariant). One unique step to developing an SDE requires the user to specify the vector of changes for each transition, i.e. a vector denoting changes in the two populations involved in a transition: the initial state (−1) and the destination states (+1). After providing these vectors, the drift vector and the diffusion matrix can be calculated, which then completely specifies an SDE model. The users are then tasked with choosing a solver and conducting a Monte Carlo simulation to estimate decision-relevant quantities. The implementation of the solver and the Monte Carlo method depends on the user’s choice of programming language. In this study, the SDE models are implemented in R and Julia, and we use an R package with a comprehensive set of SDE-related functionalities to solve SDEs.[18]

In this study, the derivation of an SDE model follows a heuristic approach. A more rigorous method seeks to prove that the form of the equation Δ**N**(*t*) satisfies (Equation 7) is a direct consequence of the following assumptions: Δ**N**(*t*) is Markovian, smooth in its argument, and has finite mean and variance. Readers seeking a more detailed derivation should consult [24]. In addition, we do not show the relationship between the Fokker-Planck equation (FPE), which governs the time evolution of the probability function of a real-valued random vector (cf. Equation 3) and an SDE. In theory, the probability function of the solution to an SDE model satisfies an FPE. Moreover, an FPE is indeed a continuous approximation of a master equation thereby establishing a theoretical linkage between a cohort model (an evolution of a deterministic vector) and its stochastic counterpart, an SDE (an evolution of a stochastic vector). This fundamental fact can be shown through the machinery of stochastic calculus which is beyond the scope of this study. For a rigorous derivation, we refer readers to [25]. In summary, we opt for a simpler approach so potential users can readily appreciate the relationship between different representations of a cohort model and apply the methods in their works.

This study focuses on continuous-time cohort models and assumes that the discrete-time transition probability matrix is embeddable to a continuous-time model. Discrete-time Markov models have been the preferred method in many applications. However, as the dynamic process of interest often involves a disease progression component, a continuous-time model would be more appropriate in most applications.[26] Nevertheless, in some cases, an individual-level model naturally follows a discrete-time process, e.g. the transitions among living arrangements for a dementia patient occur in discrete time steps.[27] However, if we consider a sufficiently large cohort of individuals, the probability of one individual transitioning in a small time interval would very likely be nonzero. Nevertheless, future studies should investigate under what conditions a typical Markov model in decision modeling is embeddable.

## 6 Conclusions

Stochastic models can serve as powerful tools for aiding decision-makers in estimating decision-relevant outcomes and quantifying the impact of uncertainty on decision-making. Cohort model provides an easy-to-implement method for modeling recurrent events over time and has been used extensively in many applications ranging from clinical decision making to policy evaluations. The stochastic differential equation method expands the functionality of cohort models in practice by providing information on the variance of the process, decreasing computational time of Monte Carlo methods, and without necessitating additional data beyond what is usually required for cohort models.

## A Derivation of SDE

This section provides more details on the heuristic approach for deriving a stochastic differential equation (SDE) model. We use the same notations and definitions as described in the main text. Let **A**(**n**, *t*) and **B**(**n**, *t*) be the drift vector and the diffusion matrix, respectively. From the definition, we note that **B**(**n**, *t*) is an *s* × *s* positive semi-definite matrix and hence its eigenvalues are non-negative. Another consequence of positive semi-definiteness is that Σ(Δ**N**(*t*)) has a unique positive semi-definite square root, i.e. we can find a matrix **S** such that 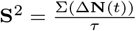. One can show that 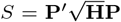, [11] where **P** is an orthogonal matrix (i.e. **PP**′ = **I** where **I** is an identity matrix) and **H** is a diagonal matrix of eigenvalues of **B**.

For a sufficiently large **N**(*t*) and using the Central Limit Theorem, Δ**N**(*t*) is approximately normal with mean **A**(**n**, *t*)*τ* and variance **B**(**n**, *t*)*τ*. Let *ξ*(*t*) be a *s*-dimensional standard Wiener process, i.e. 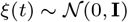.[5] Therefore we can express Δ**N**(*t*) as a function of the drift vector and the diffusion matrix, i.e. 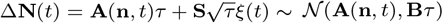. Then, the evolution of a real-valued stochastic cohort model can be represented by the following updating formula:

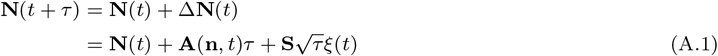

If *τ* approaches 0, then we have 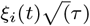 approaching *dW_i_*(*t*) at the limit (*W*(*t*) is a Wiener process) and arrive at the *Itô* SDE:

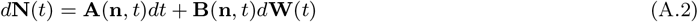

where **W**(*t*) be a *s*-dimensional Wiener process. if for all *t* and *t*_0_, the following integral equation holds:

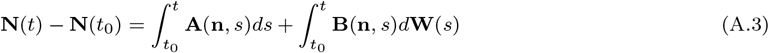

The state-wise SDE is:

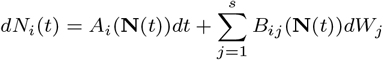

with the Ito integral equation:

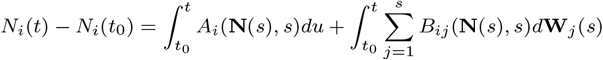

for *i* = 1, 2, …, *s*.

## Notes

This study received no financial support from any institutions or grants.

#### Summary of Updates

The revised manuscript has the following major additions/revisions: 1. The second case study has more detailed instructions which correspond to the step-by-step guidance in Section 3. 2. The detailed derivation of an SDE model is moved to the appendix to increase the accessibility and readability of the main text. 3. The section on the Fokker-Planck Equation has been removed.

